# Untangling a History of Hybridization: A Comparison of Phylogenetic Network Methods in Reconstructing Reticulate Evolution in New Zealand Cicadas

**DOI:** 10.1101/2025.03.04.641558

**Authors:** Mark Stukel, Chris Simon

## Abstract

Rapid species radiations make hybridization among species more likely. Detecting and reconstructing hybridization is therefore critical for understanding species relationships in many cases. We explored the relative performace of two phylogenetic network methods, SNaQ and PhyNEST, in evaluating the likelihood of proposed past hybridization hypotheses. We used a phylogenomic dataset of over 500 nuclear AHE genes along with mitochondrial genomes to investigate hybridization in a radiation of New Zealand cicada genera *Kikihia* and *Maoricicada*. We generated hypotheses for the existence of hybridization events based on patterns of mito-nuclear discordance and hybrid mating signals observed in a previous study. We used the D-statistic to test for the signal from hypothesized hybridization and compared SNaQ and PhyNEST results in reconstructing our hypothesized hybridization events. We found that New Zealand cicadas have an extensive history of reticulate evolution matching predictions based on patterns of mito-nuclear discordance. We also found differences in the biological plausibility of networks inferred between the two network programs, and suggest areas of improvement for network method developers.

## Introduction

Hybridization has become recognized as an important element in species radiations. Historically, hybridization among animal species was considered uncommon, which helped contribute to the development of the Biological Species Concept (Mayr 1963; Wang et al. 2020). However, research has increasingly shown that reproductive isolation between animal species is not always complete and species boundaries are fuzzy. Some examples in animals include *Heliconia* butterflies (Mallet et al. 2007), wolves and coyotes (Kays et al. 2009), monitor lizards (Pavón-Vázquez et al. 2021), rails (Coster et al. 2018), and humans (Green et al. 2010). Hybridization scenarios in species radiations include speciation with gene flow (Nosil 2008) and hybridization after secondary contact between lineages (Butlin et al. 2008). As a result, hybridization can affect the diversification dynamics of the radiation through hybrid speciation or the transfer of adaptive genes across species boundaries (Mallet et al. 2016). Methods for detecting and reconstructing hybridization events have become increasingly important for understanding how hybridization affects species radiations.

Methods for detecting hybridization search for deviations from patterns expected by incomplete lineage sorting (ILS), also called deep coalescence (Hibbins and Hahn 2022). There are three possible tree topologies for a rooted triplet or unrooted quartet of species. In the absence of ILS or hybridization, all of the gene trees within a species tree will match the species tree topology for this rooted triplet or unrooted quartet. When ILS is present, some gene trees will have one of the two alternative topologies for the rooted triplet or unrooted quartet instead of the topology agreeing with the species tree; this is caused by the lineages in these gene trees coalescing in a common ancestor of all three taxa. In the presence of ILS alone, the multi-species coalescent (MSC) predicts that the proportion of gene trees exhibiting the two alternative topologies will be equal to one another, because in the ancestral species lineage the order in which lineages coalesce is random (Pamilo and Nei 1988; Takahata 1989; Maddison 1997; Rosenberg 2002). The presence of hybridization in this rooted triplet will skew the proportion of alternative gene tree topologies because there is an alternative way that lineages in a gene tree can coalesce, which is through the hybridization event (Hahn and Hibbins 2019). The most widely-used test for detecting hybridization, the D-statistic or ABBA-BABA test, checks whether site patterns matching the alternative gene tree topologies are significantly unequal across a genome, while other methods look for unequal frequencies of gene tree topologies (Green et al. 2010; Hahn and Hibbins 2019).

To go beyond simple detection of hybridization, phylogenetic network methods are used to reconstruct the history of hybridization events among species. Some methods for constructing phylogenetic networks such as SplitsTree infer implicit networks, which visualize the patterns of conflicting phylogenies within a dataset (Huson 1998). As the history of reticulation events can be difficult to interpret from implicit networks, methods for inferring explicit networks that describe the reticulation history have become relevant (Huson and Bryant 2006). Much like tests for detecting hybridization such as the D-statistic, it is extremely important that phylogenetic network methods are able distinguish between patterns in the data caused by ILS and patterns caused by hybridization. Modern methods for inferring explicit phylogenetic networks therefore incorporate the network extension of the MSC, the network multi-species coalescent (NMSC), which takes into account the effects of both ILS and hybridization (Meng and Kubatko 2009). Various methods using the NMSC to infer phylogenetic networks have been developed. These include methods which use maximum parsimony (Yu et al. 2013), maximum likelihood, (Yu et al. 2014), maximum composite or pseudolikelihood (Yu and Nakhleh 2015; Solís-Lemus and Ané 2016; Allman et al. 2019), and Bayesian inference (Wen et al. 2016) on input gene trees, as well as methods which use maximum pseudolikelihood (Kong et al. 2024) and Bayesian inference (Wen et al. 2016; Zhang et al. 2018) on sequence data. Maximum pseudolikelihood methods, which compute likelihoods of 4-taxon quartets under the NMSC before multiplying them to find the composite likelihood of the full network, have shown promise for their speed and scalability. One such method is SNaQ (Solís-Lemus and Ané 2016), a popular method implemented in the PhyloNetworks Julia package (Solís-Lemus et al. 2017), which conducts maximum pseudolikelihood inference on quartets from gene tree input. An experimental alternative method which also uses maximum pseudolikelihood inference on quartets is PhyNEST (Kong et al. 2024), which uses site patterns from sequence data as input.

Here we compare the performance of SNaQ and PhyNEST, two maximum pseudolikelihood phylogenetic network methods which use different types of data as input, using a New Zealand (NZ) cicada dataset in which hybridization events have previously been supported by song, and mitochondrial DNA phylogenies compared to nuclear gene phylogenies (Marshall et al. 2011; Wade 2014; Stukel et al. 2024). The NZ cicada genera *Kikihia*, *Maoricicada*, and *Rhodopsalta* form a clade of around 45 species and subspecies descending from a single colonization event followed by a rapid species radiation (Arensburger et al. 2004a). The roughly thirty species of *Kikihia* are specialists on grassland, forest, evergreen shrub, and dry scrub habitats and range in altitude from sea level to subalpine (Arensburger et al. 2004a, 2004b; Marshall et al. 2008). The genus *Maoricicada* consists of 14 species, five of which are found in lowland habitats and the remaining nine forming a radiation into alpine and subalpine habitats (Buckley and Simon 2007). The three species of *Rhodopsalta* are restricted to coastal, lowland, and mid-elevation areas and have not undergone the same diversification as the other two genera (Bator et al. 2021). Previous work on these NZ cicadas has identified extensive recent hybridization among many *Kikihia* species across various hybrid zones using Sanger-sequenced loci and microsatellites (Marshall et al. 2008, 2011; Wade 2014). Using phylogenomic datasets, Stukel et al. (2024) identified strong discordance between nuclear phylogenies and mitochondrial genome phylogenies, as well as discordance among nuclear gene trees and site patterns, in both *Kikihia* and *Maoricicada* that may be the result of hybridization events. If the mito-nuclear discordance and discordance among nuclear gene trees and site patterns are the result of hybridization, then the mito-nuclear discordance results should be compatible with phylogenetic networks inferred from nuclear genes and mitochondrial genomes in these phylogenomic datasets.

Using the mito-nuclear discordance results from Stukel et al. (2024) as a basis, we tested the following hybridization hypotheses (Fig. 1).

**Figure 1:**
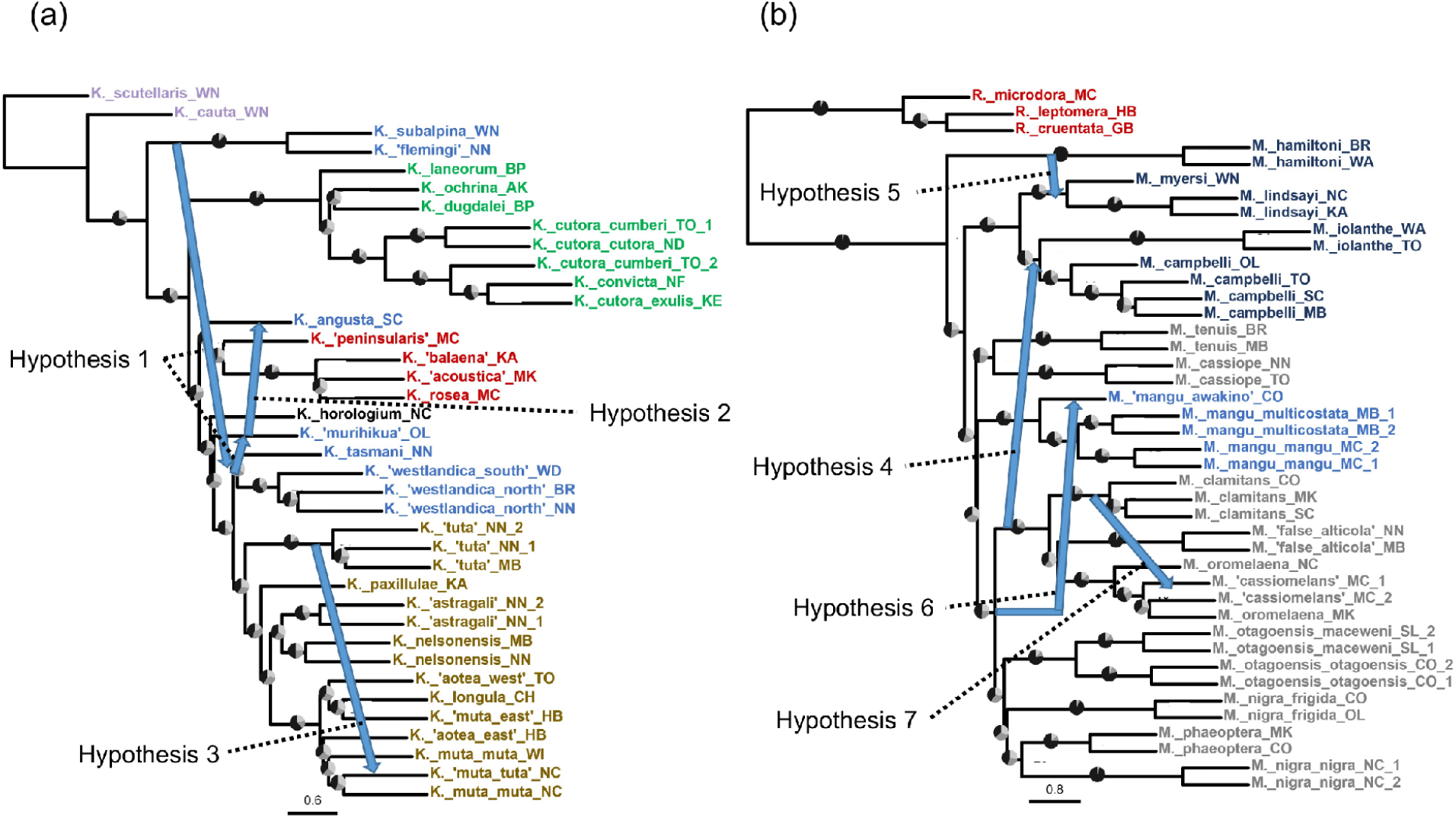
Hypothesized hybridization events. Hybridization events predicted by Hypotheses 1-7 plotted on (a): *Kikihia* ASTRAL phylogeny modified from Stukel et al. (2024) (b): *Maoricicada* ASTRAL phylogeny modified from Stukel et al. (2024).

### *Hypothesis 1:* “The Westlandica hypothesis”

The ancestor of the subalpine forest edge *Kikihia* species *K. subalpina* and *K.* “flemingi” hybridized with the ancestor of the grass species *K.* “tasmani”, *K.* “westlandica north”, and *K.* “westlandica south”, and said ancestor hybridized with the tussock grass species *K. angusta* or the dry scrub species *K.* “murihikua”. The three grass species form a clade with the other grass species in nuclear phylogenies, but in mitochondrial phylogenies all seven species form a clade nicknamed the “Westlandica Group” (Marshall et al. 2008; Stukel et al. 2024).

### *Hypothesis 2:* “The *Kikihia angusta*-‘murihikua’ hypothesis”

The tussock grass species *K. angusta* hybridized with the dry scrub species *K.* “murihikua”. Stukel et al. (2024) recovered these two species as sister to each other in mitochondrial and concatenated nuclear phylogenies, while in the coalescent nuclear phylogenies they recovered the two species as separated, suggesting hybridization as the cause of this discordance.

### *Hypothesis 3:* “The *Kikihia muta*-‘tuta’ hypothesis”

The grass species *K. muta muta* hybridized with the grass species *K.* “tuta” to form the hybrid population *K.* “muta-tuta”. Nuclear phylogenies show *K.* “muta-tuta” grouping with the two *K. m. muta* individuals in the nuclear phylogenies, but grouping with sampled *K.* “tuta” individuals in mitochondrial phylogenies, suggesting hybridization followed by mitochondrial capture (Stukel et al. 2024). Additionally, previous work using microsatellites and mitochondrial DNA suggest ongoing hybridization between *K. m. muta* and *K.* “tuta” in the range of *K.* “muta-tuta” (Wade 2014).

### *Hypothesis 4*: “The *Maoricicada campbelli* hypothesis”

The ancestor of the lowland species *M. iolanthe* and the lowland and subalpine species *M. campbelli* hybridized with the ancestor of the alpine species *M. clamitans* and *M. oromelaena*. Stukel et al. (2024) recovered *M. campbelli* and *M. iolanthe* as members of a lowland *Maoricicada* clade with *M. myersi* and *M. lindsayi* in nuclear phylogenies, but forming a clade with *M. clamitans* and *M. oromelaena* nested within the alpine radiation in the mitochondrial phylogeny. This discordance suggests hybridization followed by mitochondrial capture.

### *Hypothesis 5*: “The *Maoricicada hamiltoni* hypothesis”

The lowland species *M. hamiltoni* hybridized with the ancestor of the lowland species *M. myersi* and *M. lindsayi*. *M. hamiltoni* is recovered as sister to all other *Maoricicada* in nuclear phylogenies, but recovered as sister to *M. myersi* and *M. lindsayi* in the mitochondrial genome phylogeny, suggesting hybridization followed by mitochondrial capture (Stukel et al. 2024).

### *Hypothesis 6*: “The *Maoricicada mangu* hypothesis”

The subalpine species *M. mangu* hybridized with an unsampled “ghost” lineage of alpine *Maoricicada*, forming a hybrid population *M.* “mangu awakino” at the Awakino Ski Field locality. Stukel et al. (2024) recover this individual as grouping with its conspecifics *M. m. mangu* and *M. m. multicostata* in nuclear phylogenies, but recover this individual elsewhere in the alpine radiation with uncertain placement in the mitochondrial phylogeny. This discordance suggests hybridization with an unknown species followed by mitochondrial capture.

### *Hypothesis 7:* “The *Maoricicada* ‘cassiomelans’ hypothesis”

The individuals labeled *M.* “cassiomelans” are hybrids involving the alpine species *M. oromelaena*, *M. clamitans*, and *M. cassiope*. The *M.* “cassiomelans” individuals, which have very similar morphology to *M. oromelaena*, were observed to have unusual song characteristics that differentiated them from *M. oromelaena*. The original collectors of the *M.* “cassiomelans” individuals noted the similarity of the songs to *M. clamitans* and *M. cassiope*, suggesting hybrid origin with these species. However, there is no mitonuclear discordance data to suggest this hybridization event.

## Methods

Our data for this study consisted of the sequence alignments for the nuclear Anchored Hybrid Enrichment (AHE) loci and mitochondrial genomes produced by Stukel et al. (2024), obtained from Dryad (https://doi.org/10.5061/dryad.t1g1jwt7v). The data were divided into a *Kikihia* dataset and a *Maoricicada* + *Rhodopsalta* dataset. We ran analyses using SNaQ and PhyNEST. To test each hybridization hypothesis, we chose a subset of taxa that included the hypothesized hybrids and parent lineages while excluding taxa uninvolved with the hybridization and unneeded for phylogenetic placement. This allowed us to reduce the number of taxa in each analysis for computational tractability. Current phylogenetic network methods are restricted to inferring level-1 networks, which are networks in which no two reticulation events share a network edge (Rosselló and Valiente 2009). Because Hypothesis 1 involves multiple hybridization events with the same taxa which might not be resolvable in a level-1 network, we created four overlapping taxon subsets for inference instead of just one. These four taxon subsets would allow us to potentially isolate individual hybridization events that might otherwise interfere with one another, provided they are not events between sister taxa. Additionally, in our preliminary investigations of Hypothesis 4 we found potentially conflicting hybridization events, so we decided to create two taxon subsets for inference, one focusing on hybridization with alpine species and one focusing on hybridization with lowland species. To more easily compare the results of SNaQ and PhyNEST, we used the same taxon subsets for analysis with each program. The list of taxon subsets are found in Table 1.

**Table 1:** Taxa used in phylogenetic network taxon sets. See table in Stukel et al. (2024) for full taxon information.

| Taxon set | Hypothesis for testing | Taxa included |
| --- | --- | --- |
| flemingi-angusta-murihikua | Hypothesis 1 | K._angusta_08.NZ.SC.IDN.12; K._flemingi_11.NZ.NN.LGU.02; K._murihikua_08.NZ.OL.RSC.01; K._muta_muta_12.NZ.NC.BAL.01; K._scutellaris_14.NZ.WN.NEV.03 |
| flemingi-westlandica | Hypothesis 1 | K._flemingi_11.NZ.NN.LGU.02; K._muta_muta_12.NZ.NC.BAL.01; K._scutellaris_14.NZ.WN.NEV.03; K._tasmani_02.NZ.NN.COR.21; K._westlandica_north_12.NZ.BR.MMK.17; K._westlandica_south_11.NZ.WD.OKT.33 |
| westlandica-angusta-murihikua | Hypothesis 1 | K._angusta_08.NZ.SC.IDN.12; K._murihikua_08.NZ.OL.RSC.01; K._muta_muta_12.NZ.NC.BAL.01; K._scutellaris_14.NZ.WN.NEV.03; K._tasmani_02.NZ.NN.COR.21; K._westlandica_north_12.NZ.BR.MMK.17; K._westlandica_south_11.NZ.WD.OKT.33 |
| full-westlandica | Hypothesis 1 | K._angusta_08.NZ.SC.IDN.12; K._flemingi_11.NZ.NN.LGU.02; K._murihikua_08.NZ.OL.RSC.01; K._muta_muta_12.NZ.NC.BAL.01; K._scutellaris_14.NZ.WN.NEV.03; K._tasmani_02.NZ.NN.COR.21; K._westlandica_north_12.NZ.BR.MMK.17; K._westlandica_south_11.NZ.WD.OKT.33 |
| angusta-murihikua | Hypothesis 2 | K._angusta_08.NZ.SC.IDN.12; K._murihikua_08.NZ.OL.RSC.01; K._muta_muta_12.NZ.NC.BAL.01; K._rosea_11.NZ.MC.CHZ.02; K._scutellaris_14.NZ.WN.NEV.03 |
| muta-tuta | Hypothesis 3 | K._scutellaris_14.NZ.WN.NEV.03; K._tuta_05.NZ.MB.TFL.03; K._muta_muta_12.NZ.NC.BAL.01; K._muta_tuta_12.NZ.NC.WAI.14; K._paxillulae_18.NZ.KA.KAI.3 |
| campbelli-alpine | Hypothesis 4 | M._campbelli_02.NZ.OL.FRL.04; M._clamitans_02.NZ.CO.AWT.14; M._iolanthe_02.NZ.WA.BUL.02; M._mangu_mangu_10.NZ.MC.HUT.06; M._phaeoptera_02.NZ.CO.AWT.13; R._leptomera_06.NZ.HB.POR.12 |
| campbelli-lowland | Hypothesis 4 | M._campbelli_02.NZ.OL.FRL.04; M._iolanthe_02.NZ.WA.BUL.02; M._lindsayi_06.NZ.KA.CAM.04; M._myersi_02.NZ.WN.ORO.03; R._leptomera_06.NZ.HB.POR.12 |
| hamiltoni | Hypothesis 5 | M._cassiope_14.NZ.NN.STE.04; M._hamiltoni_02.NZ.BR.MRV.02; M._lindsayi_06.NZ.KA.CAM.04; M._mangu_mangu_10.NZ.MC.HUT.06; M._myersi_02.NZ.WN.ORO.03; R._leptomera_06.NZ.HB.POR.12 |
| awakino | Hypothesis 6 | M._false_alticola_14.NZ.MB.RST.01; M._mangu_awakino_17.NZ.CO.AWC.03; M._mangu_mangu_10.NZ.MC.HUT.06; M._phaeoptera_02.NZ.CO.AWT.13; R._leptomera_06.NZ.HB.POR.12 |
| cassiomelans | Hypothesis 7 | M._cassiomelans_12.NZ.MC.HUT.03; M._cassiope_14.NZ.NN.STE.04; M._clamitans_02.NZ.CO.AWT.14; M._oromelaena_02.NZ.NC.TBS.01; R._leptomera_06.NZ.HB.POR.12 |

### SNaQ analyses (gene trees as input)

We used the TICR pipeline (Stenz et al. 2015) to prepare gene tree input data for SNaQ as this works to account for gene tree uncertainty. This pipeline uses alignments input into MrBayes 3.2.7 (Ronquist et al. 2012) to estimate posterior distributions of gene trees. It then uses those gene tree posteriors as input for BUCKy (Larget et al. 2010) to generate a table of quartet concordance factors (CF) with credible intervals for each 4-taxon subset. This table of quartet CFs and credible intervals is then used as input for SNaQ.

For the MrBayes runs on the AHE nuclear loci, we used GTR+I+G for the substitution model treating each locus as a single partition. For the runs on the mitochondrial genome trees, we used GTR+I+G for the substitution model and used the following data partitions: combined first and second codon positions for all protein-coding genes, third codon positions for all protein-coding genes, 12S rRNA, 16S rRNA, all tRNAs combined. We used the following settings for the MCMC on all the nuclear and mitochondrial alignments following the guidelines in the TICR pipeline: 3 independent runs, 1 million generations per run, 3 chains (1 cold and 2 heated), heated chain temperature of 0.4, burn-in fraction of 0.25, swap frequency of every 10 generations, and sample frequency of every 50 generations. We expected that the mitochondrial genome trees would be the slowest to converge since they used the most complicated model; convergence was assessed for the mitochondrial genome trees using the *sump* command.

Following the TICR pipeline, we summarized the MrBayes tree posteriors for all three runs for each nuclear locus and mitochondrial genome alignment and generated quartet CF tables and credible intervals with BUCKy. We ran BUCKy using the default settings outlined in the TICR pipeline. We pruned the resulting concordance factor (CF) tables into smaller tables, each consisting of the taxa in a particular taxon subset for hypothesis testing.

For our initial network searches, we performed SNaQ runs on each taxon subset with a range of maximum number of hybridization events (hmax) using the CF tables as input and the ASTRAL tree topology from Stukel et al. (2024) as the starting topology. We plotted the network log pseudolikelihood for each hmax value and selected as our candidate network the network with the hmax value where the log pseudolikelihood transitioned from steep improvement to slow, linear improvement. Since SNaQ does not infer rooted networks, there is a possibility that the candidate network will have a hybridization edge that conflicts with the root placement (see Discussion below). In addition to the best network found in a run, SNaQ provides a list of alternative networks with different hybrid edge placements for the same reticulation cycle along with their pseudolikelihoods. This allowed us to choose a version of the inferred network with hybridization edges compatible with the outgroup if the inferred network conflicted with the outgroup taxon. To account for gene tree uncertainty in the estimation of our candidate networks, we used the SNaQ *bootsnaq* function to generate 100 bootstrap replicates for each candidate network using the CF 95% credible intervals as input. These bootstrap networks were used to annotate the candidate network, with the exception of two runs in which the candidate network was not recovered in any of the bootstrap networks. In those cases, we selected an alternative network consistent with the tested hypothesis from the list of bootstrap networks to annotate.

### PhyNEST (site patterns as input)

We conducted PhyNEST analyses on two datasets for each taxon set: nuclear AHE loci only and nuclear AHE loci + mitochondrial genomes. Because the mitochondrial genome has a higher proportion of informative sites compared to the nuclear AHE genes, signal from the mitochondrial genome might overwhelm any signal from the nuclear genes. To examine whether the mitochondrial genome had a disproportionate effect on the PhyNEST results, we used alignments that included the nuclear AHE genes alone as well as alignments that included both nuclear AHE genes and mitochondrial genomes. Since the mitochondrial genome acts as a single locus and SNaQ treats all loci equally, we did not need to adjust for the disproportionate effect of the mitochondrial genome in our SNaQ analyses. We performed network searches on each taxon subset with a range of hmax values using both the hill-climbing and simulated annealing search strategies. PhyNEST requires an outgroup taxon to be designated in order to return a rooted network. All the *Kikihia* taxon set analyses were rooted using *K. scutellaris*, which is the species known to be sister to all other *Kikihia* based on previous phylogenetic analyses (Marshall et al. 2011 and references therein), while all the *Maoricicada* taxon set analyses were rooted using *Rhodopsalta leptomera*.

The hill-climbing and simulated annealing analyses were all run using default settings in PhyNEST, with the exception of the *cons* and *alph* parameters for the simulated annealing analyses, which were each set to 0.25. As with the SNaQ analyses, we selected the network with the hmax value where the log pseudolikelihood transitioned from steep improvement to slow, linear improvement. A few analyses had massive improvements in log pseudolikelihood on the highest hmax run compared to the others. After reviewing the networks for these runs, we found hybridization edges with near zero inheritance probabilities, so based on the advice in the PhyNEST documentation we selected the networks with the next highest hmax value (the value before the massive improvement in log pseudolikelihood). PhyNEST does not currently have a built-in method to assess uncertainty in the network inference e.g. bootstrapping. In principle, bootstrap replicates could be generated manually, but we did not do so. We also used the *Dstat* function in PhyNEST to perform ABBA-BABA tests on all sets of four taxa to test hybridization hypotheses (Durand et al. 2011). We performed the test on the PhyNEST datasets with and without mitochondrial DNA. We used a Bonferroni correction to account for multiple comparisons when calculating the significance value for the ABBA-BABA tests. The data and scripts for this study are deposited in Dryad and Zenodo (http://datadryad.org/stash/share/MSubfB-NR9rOzzKWJMbnbtcK-Y3mX5YWmOZDTcGp0Jo).

## Results

The results from SNaQ and PhyNEST were often incongruent with one another (Fig. 2-5). Most SNaQ networks had high bootstrap support. The inclusion vs. exclusion of mitochondrial DNA had a limited effect on the PhyNEST results. The results for the different hybridization hypotheses are presented below. The term “major hybrid edge” refers to a hybridization edge with inheritance probability greater than 0.5, while “minor hybrid edge” refers to hybridization edges with inheritance probability less than 0.5, where the inheritance probability is the proportion of the genome a hybrid inherited from each parent (Solís-Lemus et al. 2017). Similarly, the term “major-tree” refers to the tree represented by the major hybrid edges of a network, excluding the minor hybrid edges.

**Figure 2:**
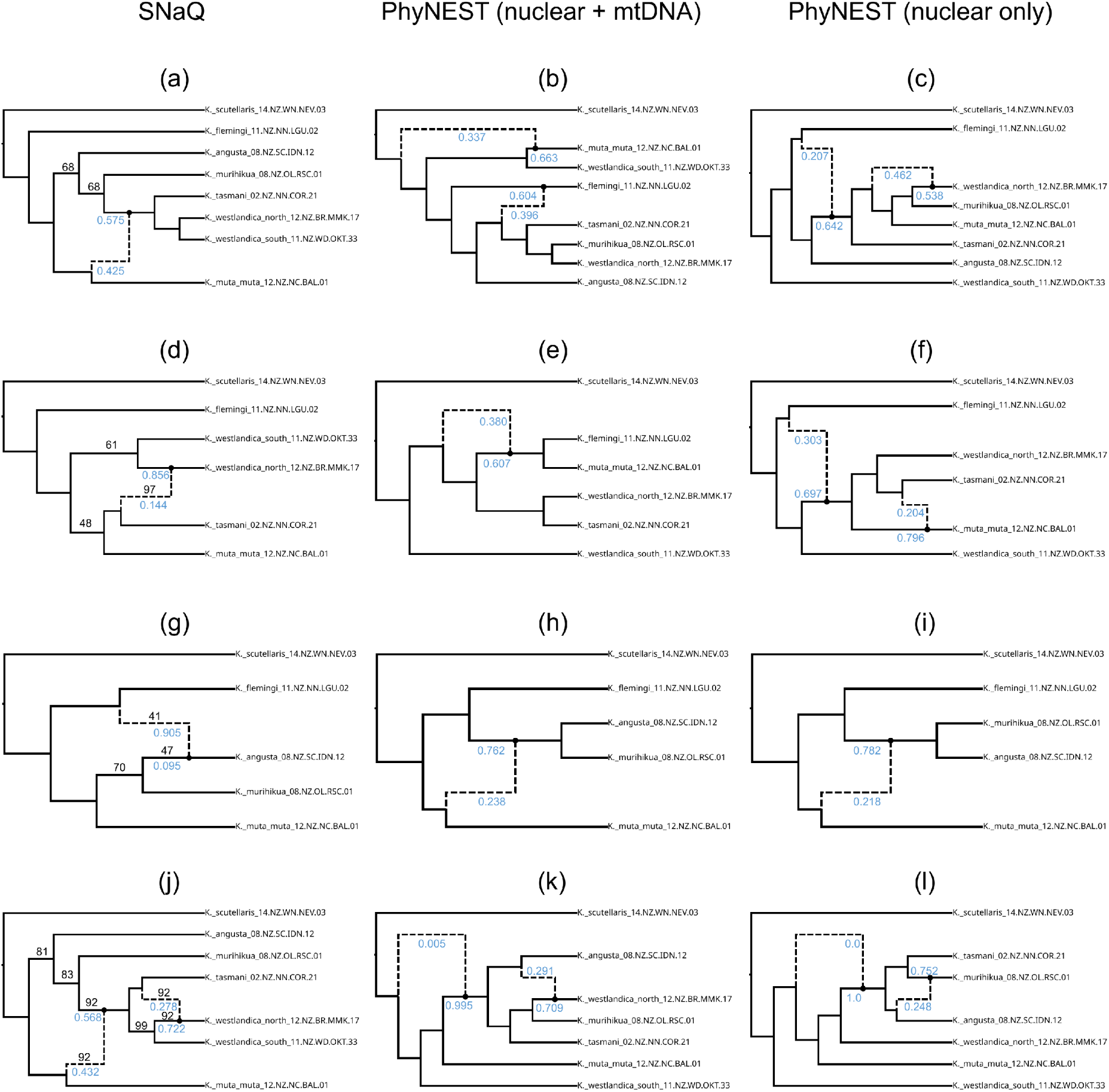
Phylogenetic network results for Hypothesis 1, separated out by taxon subset and dataset. (a)-(c): Results from the “full westlandica” taxon set. (d)-(f): Results from the “flemingi-westlandica” taxon set. (g)-(i): Results from the “flemingi-angusta-murihikua” taxon set. (j)-(l): Results from the “angusta-murihikua-westlandica” taxon set. Inheritance probabilities for hybrid edges are displayed in blue. SNaQ bootstrap proportions below 100 displayed in black on SNaQ networks only.

### *Hypothesis 1: Kikihia* “westlandica group” species

The SNaQ networks across the four taxon subsets for this hypothesis were all congruent with one another and showed similar major-tree topologies to previous nuclear phylogenomic trees (Stukel et al. 2024). These networks display hybridization events between *Kikihia muta muta* and the ancestor of *K.* “tasmani”, *K.* “westlandica north”, and *K.* “westlandica south” (Fig. 2 a, j) and between *K.* “tasmani” and *K.* “westlandica north” (Fig. 2 g, j), both with high SNaQ bootstrap support. They also display hybridization between *K.* “flemingi” and *K. angusta*, albeit with low bootstrap support (Fig. 2 d). These hybridization events are consistent with those predicted for this hypothesis.

In contrast, the PhyNEST networks across the four taxon subsets were for the most part not congruent with one another or with the SNaQ networks. They also displayed highly discordant major-tree topologies compared to previously published nuclear phylogenomic trees (Stukel et al. 2024). *K.* “westlandica south” was consistently inferred in an unusual position not sister to *K.* “westlandica north” or *K.* “tasmani” (Fig. 2 b-c, e-f, k-o). As the exception, for one taxon subset PhyNEST did infer networks that were congruent with the corresponding SNaQ network, but with the major and minor hybrid edges swapped. For this taxon subset, both the nuclear + mitochondrial DNA dataset and the nuclear-only dataset inferred a hybridization event between *K. muta muta* and the ancestor of *K. angusta* and *K.* “murihikua” (Fig. 2 h-i).

The D-statistic results showed evidence of hybridization between *K. subalpina/*“flemingi” and *K.* “tasmani”/“westlandica north”/“westlandica south” for both datasets, with and without mitochondrial DNA (Tables S1-2). They also detected hybridization between *K. subalpina/*“flemingi” and *K. angusta*/“murihikua” (Tables S1-2). The D-statistic results are consistent with the predicted hybridization events for this hypothesis.

### *Hypothesis 2: Kikihia angusta* and K. “murihikua”

The SNaQ and PhyNEST networks were not congruent with one another. The SNaQ network inferred hybridization between *K. muta muta* and *K. scutellaris* with low bootstrap support (Fig. 3a). The PhyNEST network inferred from nuclear and mitochondrial DNA combined displays hybridization between an ancestor of *K. angusta* and *K.* “murihikua” and *K. rosea* (Fig. 3b), while the network using nuclear DNA alone displays *K. angusta* hybridizing with *K.* “murihikua” while sister to *K. muta muta* and *K. rosea* (Fig. 3c). The hybridization inferred in the SNaQ network is not consistent with the predictions of this hypothesis, but both PhyNEST methods are consistent with the predictions.

**Figure 3:**
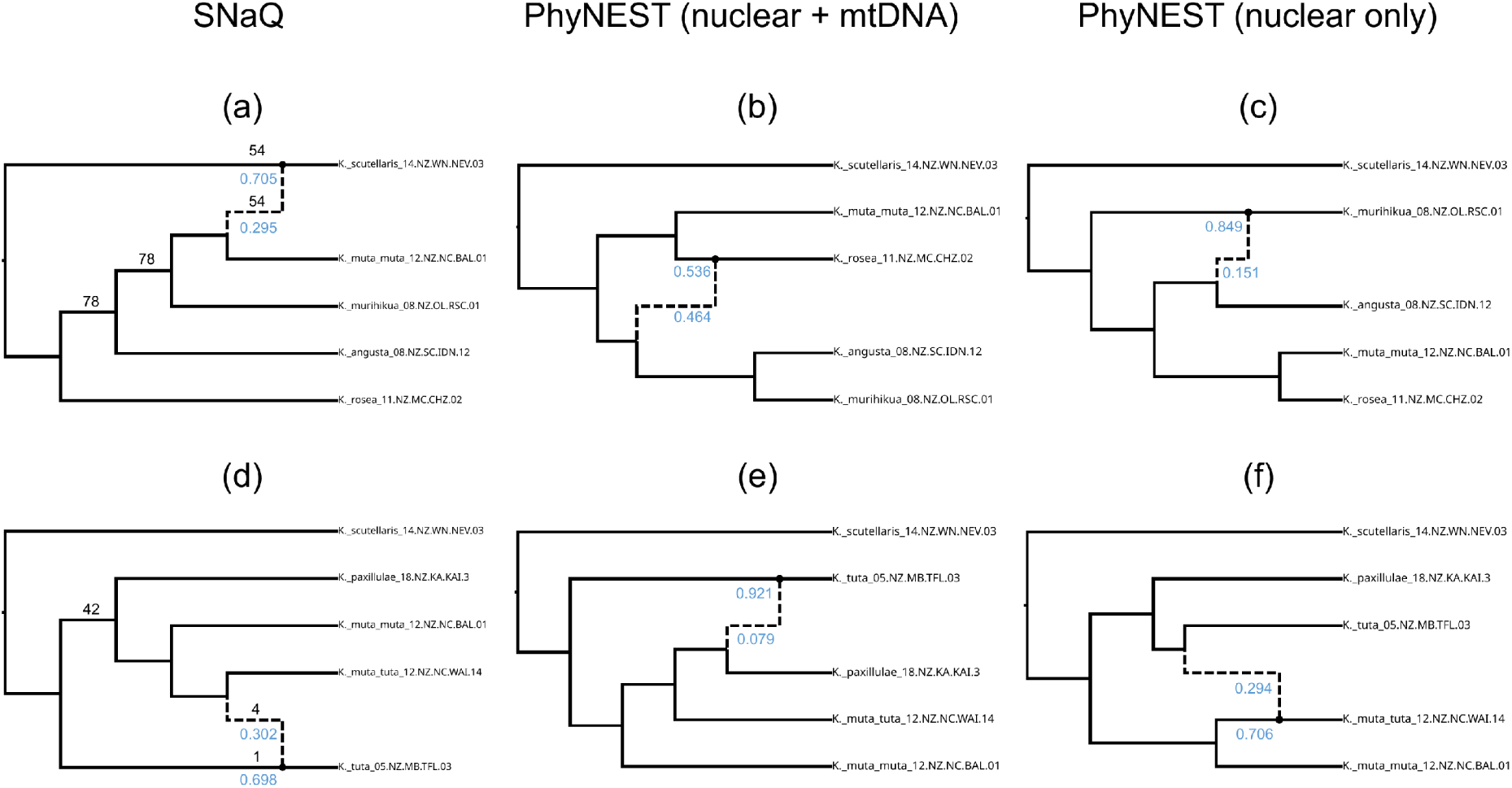
Phylogenetic network results for Hypotheses 2 and 3, separated out by dataset. (a)-(c): Results for **Hypothesis 2,** hybridization involving *Kikihia angusta* and *K.* “murihikua”. (d)-(f): Results for **Hypothesis 3,** hybrid origin of *K.* “muta-tuta”. Inheritance probabilities for hybrid edges are displayed in blue. SNaQ bootstrap proportions below 100 displayed in black on SNaQ networks only.

The D-statistic results show evidence of gene flow between *K.* “murihikua” and *K. muta muta*, between *K.* “murihikua” and *K. angusta*, between *K. angusta* and *K. rosea*, and between *K.* “murihikua” and *K. rosea* (Tables S1-2), consistent with the predictions for this hypothesis.

### *Hypothesis 3: Kikihia* “muta-tuta”

The best network inferred by SNaQ for this taxon set had hybridization edges that were incompatible with the *K. scutellaris* outgroup. Since SNaQ also provides a list of alternative networks which have the same reticulation cycle as the best network, we chose a new candidate network from this list with compatible hybridization edges and a similar pseudolikelihood score to the original best network to be our candidate network. When we conducted bootstrapping in SNaQ, this network was not found among any of the bootstrap replicates. Therefore, to test our hypothesis of hybridization between *K. muta muta* and *K.* “tuta” to produce *K.* “muta-tuta”, we selected a bootstrap network showing a hybridization event between *K.* “muta-tuta” and *K.* “tuta” to annotate. This hybridization event had very low bootstrap support (Fig. 3d). The PhyNEST network inferred from nuclear and mitochondrial DNA combined displays hybridization between *K. paxillulae* and *K.* “tuta” (Fig. 3e), while the network inferred from nuclear DNA alone displays hybridization between *K.* “tuta” and *K.* “muta-tuta” (Fig. 3f). Due to the low bootstrap support, SNaQ results are not consistent with the predicted hybridization events for this hypothesis, while the PhyNEST results are partially consistent.

The D-statistic results show evidence of hybridization between *K.* “muta-tuta” and *K.* “tuta”, consistent with the mito-nuclear discordance data and the predictions of this hypothesis (Tables S1-2).

### Hypothesis 4: Maoricicada campbelli *and* M. iolanthe

The SNaQ networks from the two taxon subsets show *M. campbelli* and *M. iolanthe* involved in hybridization with both alpine and lowland species. The taxon set focusing on alpine species was inferred to have a hybridization event between *M. clamitans* and *M. campbelli* (Fig. 4a), while the taxon set focusing on lowland species was inferred to have hybridization between *M. myersi* and *M. iolanthe* (Fig. 4d), both with bootstrap support above 99%. The major-tree topologies of these networks were also consistent with previously published nuclear phylogenies (Stukel et al. 2024).

**Figure 4:**
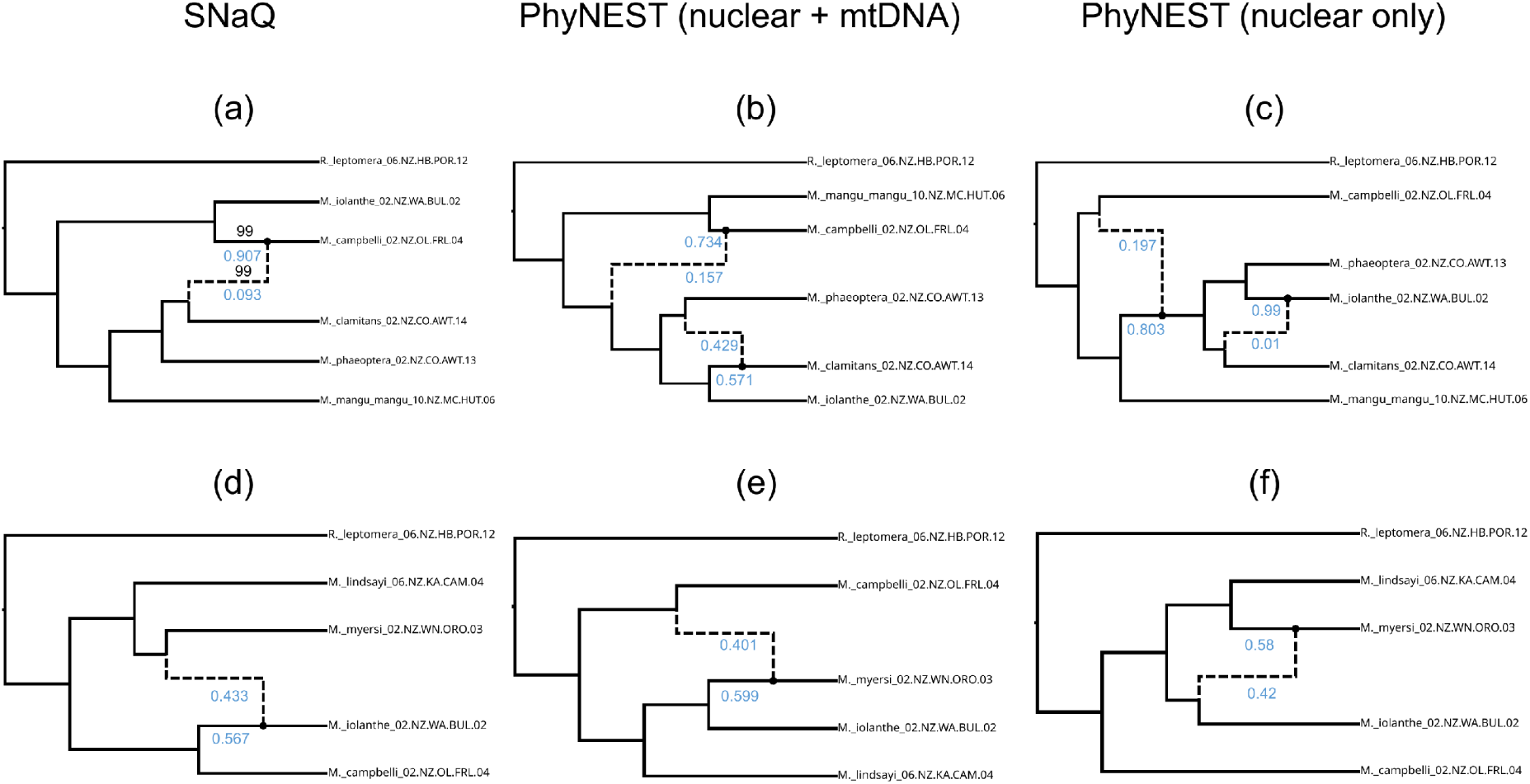
Phylogenetic network results for Hypothesis 4, separated out by taxon subset and dataset. (a)-(c): Results for the “alpine” taxon set. (d)-(f): Results for the “lowland” taxon set. Inheritance probabilities for hybrid edges are displayed in blue. SNaQ bootstrap proportions below 100 displayed in black on SNaQ networks only.

The PhyNEST networks disagreed substantially between the nuclear + mitochondrial DNA analysis and the nuclear DNA only analysis for one of the taxon sets, but were in agreement for the other taxon set. For the alpine species taxon set, PhyNEST inferred two hybridization events for both the nuclear only and nuclear + mtDNA datasets, but neither the hybrid edges nor the major-tree topologies agreed (Fig. 4b-c). The major-tree topologies were also inconsistent with previously published phylogenies (Stukel et al. 2024). In contrast, the two PhyNEST datasets on the lowland species taxon set were similar to each other and to the SNaQ network. The analysis with both nuclear and mitochondrial DNA recovered a hybridization event between *M. campbelli* and *M. myersi* (Fig. 4e), while the analysis with only nuclear DNA recovered a hybridization event between *M. iolanthe* and *M. myersi* (Fig. 4f). These two networks on the lowland species set have identical topologies except for the hybrid edges: the major hybrid edge in the nuclear only dataset was the minor edge in the nuclear + mtDNA dataset and vice versa.

The D-statistic results show evidence of gene flow between *M. campbelli* and *M. clamitans* for some *M. campbelli* individuals, and gene flow between *M. iolanthe* and *M. clamitans* for other individuals (Table S2). They also show evidence of gene flow between *M. myersi* and *M. iolanthe*, which is consistent with the SNaQ networks and the PhyNEST networks on the lowland species.

### *Hypothesis 5*: Maoricicada hamiltoni

The SNaQ and PhyNEST results were not congruent with one another. SNaQ inferred a network with a single hybridization event between *M. hamiltoni* and *M. cassiope* with medium SNaQ bootstrap support (Fig. 5a). The SNaQ network’s major-tree topology was also consistent with previously published nuclear phylogenies (Stukel et al. 2024). In contrast, the PhyNEST analyses on the two datasets, nuclear loci only and nuclear + mtDNA, inferred networks with two hybridization events that were incongruent with each other and that were incongruent with previous nuclear phylogenies (Fig. 5b-c) (Stukel et al. 2024).

**Figure 5:**
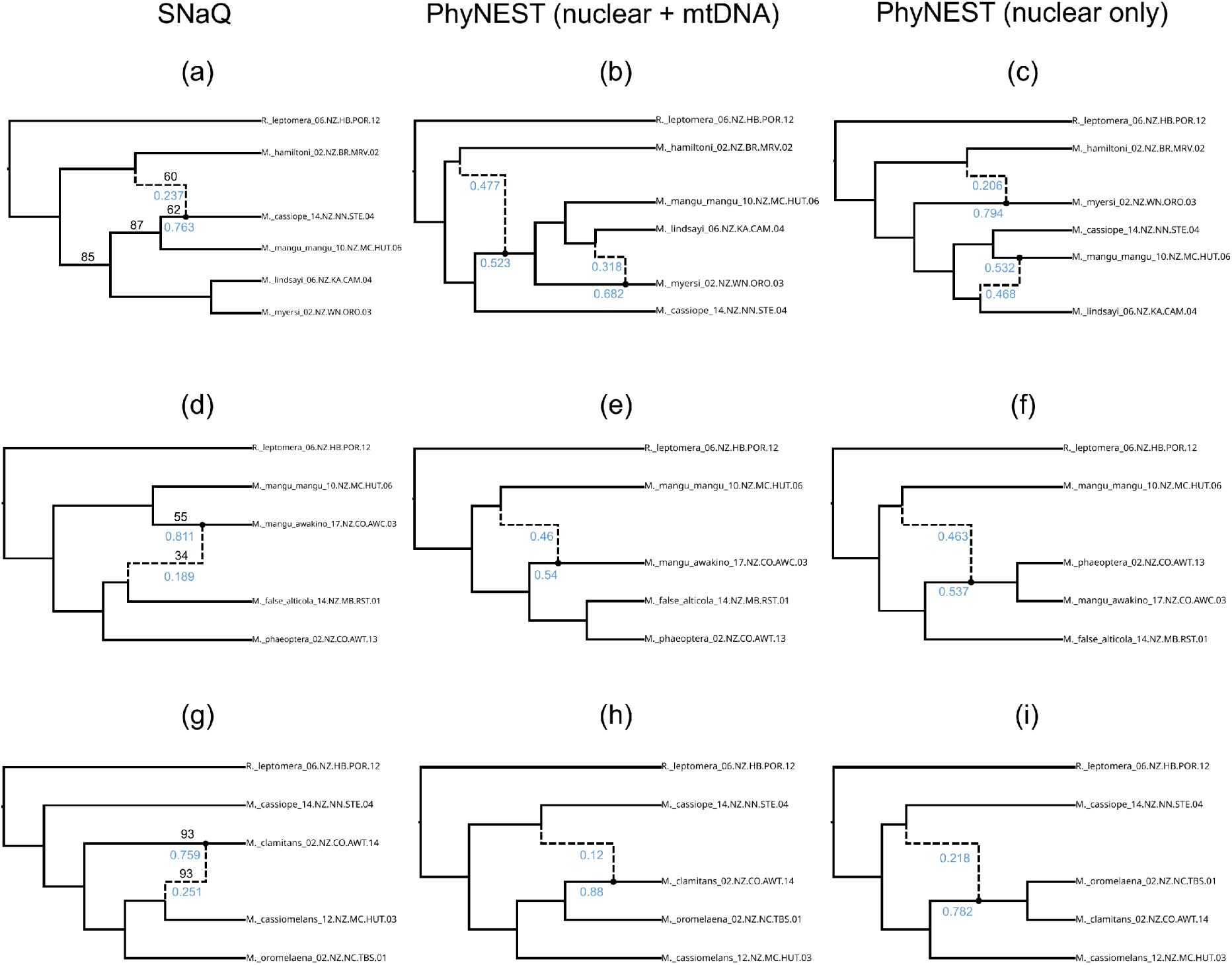
Phylogenetic network results for Hypotheses 5-7, separated out by dataset. (a)-(c): Results for Hypothesis 5, hybridization involving *Maoricicada hamiltoni*. (d)-(f): Results for Hypothesis 6, hybrid origin of *M.* “mangu awakino” population. (g)-(i): Results for Hypothesis 7, hybrid origin of *M.* “cassiomelans”. Inheritance probabilities for hybrid edges are displayed in blue. SNaQ bootstrap proportions below 100 displayed in black on SNaQ networks only.

The D-statistic results show evidence of gene flow between *M. hamiltoni* and *M. cassiope*, as well as between *M. hamiltoni* and *M. myersi*/*M. lindsayi* (Table S2).

### *Hypothesis 6: Maoricicada mangu* ghost lineage

The SNaQ and PhyNEST results were largely congruent with one another. As the original candidate SNaQ network inferred a hybridization event with zero SNaQ bootstrap support, we selected a bootstrap network showing a hybridization event between *M.* “false alticola” and *M.* “mangu awakino”, with *M.* “mangu awakino” otherwise sister to *M. mangu mangu*. This hybrid edge was recovered with low SNaQ bootstrap support, however (Fig. 5d). The PhyNEST network inferred from nuclear and mitochondrial DNA combined recovers a hybridization event between *M. mangu mangu* and *M.* “mangu awakino” (Fig. 5e). The major-tree topology of this network shows *M.* “mangu awakino” as sister to a clade containing *M. phaeoptera* and *M.* “false alticola”. The PhyNEST network inferred from only nuclear DNA recovered a hybridization event between *M. mangu mangu* and the ancestor of *M. phaeoptera* and *M.* “mangu awakino” (Fig. 5f).

The D-statistic results do not show evidence of gene flow between *M.* “false alticola” and *M.* “mangu awakino”, but they do show evidence of gene flow between *M. phaeoptera* and *M.* “mangu awakino” (Table S2).

### *Hypothesis 7: Maoricicada* “cassiomelans”

The SNaQ and PhyNEST results were slightly incongruent with one another. SNaQ inferred a network with a hybridization event between *M.* “cassiomelans” and *M. clamitans* with high SNaQ bootstrap support (Fig. 5g). In contrast, neither of the two PhyNEST networks inferred hybridization involving *M.* “cassiomelans”. The network inferred from nuclear and mitochondrial DNA combined showed hybridization between *M. cassiope* and *M. clamitans* (Fig. 5h), while the network inferred from nuclear DNA alone found hybridization between *M. cassiope* and the ancestor of *M. clamitans* and *M. oromelaena* (Fig. 5i). The major-tree topologies of both PhyNEST networks recovered *M. clamitans* and *M. oromelaena* as sister to each other, with *M.* “cassiomelans” sister to them.

The D-statistic results show evidence of gene flow between *M. clamitans* and *M.* “cassiomelans” (Table S2).

## Discussion

The results presented here provide an interesting comparison of the agreement among different analysis methods on the presence and direction of hybridization. In general, the results from SNaQ and the D-statistic tests were more consistent with the predictions of hypotheses 1-7 than were the results from PhyNEST. SNaQ supported the *M. campbelli* hypothesis, the *M. mangu* hypothesis, and the *M.* “cassiomelans” hypothesis (hypotheses 4, 6, and 7), and partially supported the Westlandica hypothesis and the *M. hamiltoni* hypothesis (hypotheses 1 and 5).

PhyNEST supported the *K. angusta-*“murihikua” hypothesis, the *K.* “muta-tuta” hypothesis, and the *M. mangu* hypothesis (hypotheses 2, 3, and 6), with partial support for the Westlandica hypothesis and the *M. campbelli* hypothesis (hypotheses 1 and 4). For all PhyNEST results, support for hypotheses was either found by both datasets, or by the nuclear-only PhyNEST dataset alone. The D-statistic results supported all hypotheses, but as detailed below they may need to be treated with some caution.

### Limitations of analysis methods

Phylogenetic network methods and hybridization tests have several limitations. As described above, current phylogenetic network methods are restricted to level-1 networks, which are networks in which no two reticulation events share a network edge (Rosselló and Valiente 2009). This complicates reconstruction of hybridization events when there are multiple rounds of hybridization or introgression, because if the same taxa are involved in multiple hybridization events the true reticulation network will not be level-1. Hypotheses 1 and 4 (Westlandica and *M. campbelli*) were split into several taxon sets in an attempt to get around this limitation, since both involve scenarios of closely-spaced hybridization events involving the same taxa. The different taxon sets for these hypotheses do show different hybridization events, but it appears that it is due to involved taxa being present or absent, rather than because the hybridization events would have interfered with one another.

Other limitations of phylogenetic networks are the root placement and direction of gene flow. While gene-tree summary methods can be affected by poorly-rooted gene trees (Mai et al. 2017), SNaQ avoids this problem by using unrooted quartets to infer a semi-directed unrooted network, which can later be rooted by the user (Solís-Lemus and Ané 2016). Since there is no indication of ancestor-descendant relationships in an unrooted network, SNaQ compares multiple hybrid positions on each reticulation cycle (Solís-Lemus and Ané 2016). However, this can sometimes result in a final network with reticulations that are incompatible with the known outgroup. To rectify this, SNaQ returns a list of alternative networks with different placements of hybrid edges on the same reticulation cycle along with their pseudolikelihood scores, so that the user can select an alternative network with a similar pseudolikelihood score that is compatible with the outgroup. For one of the taxon sets (*K.* “muta-tuta”), the network originally inferred by SNaQ was incompatible with the *K. scutellaris* outgroup, necessitating the selection of an alternative network. SNaQ is supposed to distinguish between networks with identical reticulation cycles but different hybrid edges (Solís-Lemus and Ané 2016), but if the program can produce networks with hybridization directions incompatible with known outgroups, it is conceivable that even a candidate network compatible with a known outgroup might have incorrect hybridization directions. In these cases, the judgment of the researcher on what networks are biologically plausible may be necessary.

Another analysis limitation is that PhyNEST as currently implemented may be unreliable due to the way it treats missing data in an alignment. Unlike SVDQuartets, PhyNEST currently does not account for gaps, ambiguities, or missing data in the sequence alignment and instead treats them as additional characters (Chifman and Kubatko 2014; Sungsik Kong, personal communication). Because PhyNEST uses a concatenated alignment as input, any genes that are missing some taxa or have incomplete sequence may affect the network inference, since taxa with more missing data will be treated as if they have higher sequence divergence from the other taxa than they actually do. In most cases where SNaQ and PhyNEST disagree in their network topologies, PhyNEST produces major-tree topologies that are very different from the species tree topologies presented in Stukel et al. (2024), which were inferred from the same AHE sequence data. Some of the differences in topology may be due to the different sources of data input (gene trees vs. sequence alignments/site patterns) between the two programs, but species trees inferred using those two sources of data (ASTRAL vs. SVDQuartets) did not have nearly as many differences in topology (Stukel et al. 2024). The *Kikihia* taxa chosen for our taxon subsets were recovered for 493-509 AHE nuclear genes, while the taxa in the *Maoricicada* taxon sets were recovered for 494-499 AHE nuclear genes, indicating that there was some missing data for the nuclear genes. The mitochondrial genomes were far more fragmentary for some taxa compared to others, with those for the *Kikihia* taxa ranging from 6,365 to 14,156 bp and those for the taxa in the *Maoricicada* taxon sets ranging from 9,962 to 14,068 bp. The wider range of missing data in the mitochondrial genomes may also explain the slightly worse topologies found with PhyNEST when mitochondrial DNA was included. We do not think missing data is as big of a problem for our SNaQ analyses as for our PhyNEST analyses for several reasons. First, SNaQ uses gene tree topologies as input, moving the problem of how to treat missing data to the gene tree inference method. We used MrBayes to generate the input gene trees for SNaQ, and

MrBayes excludes missing data from phylogenetic inference provided they are coded correctly in the NEXUS file (Ronquist et al. 2012). SNaQ also only calculates quartet concordance factors (CFs) from genes that have all four taxa in a quartet, regardless of whether it is calculating concordance factors from gene trees directly or using the concordance factors with credible intervals calculated by BUCKy through the TICR pipeline (Stenz et al. 2015; Solís-Lemus and Ané 2016). The PhyNEST developers are aware of the potential problem posed by missing data and are testing strategies to accommodate it in a future release (Sungsik Kong, personal communication).

Another limitation of PhyNEST as compared to SNaQ is the current lack of a measure of network branch supports. As with species tree methods based on gene tree topologies, SNaQ is sensitive to gene tree estimation error (Gatesy and Springer 2014; Roch and Warnow 2015).

While one way to reduce gene tree estimation error is to collapse poorly-supported gene tree branches (Simmons and Gatesy 2021), SNaQ can account for gene tree estimation error by first calculating credible intervals for quartet CFs from Bayesian gene tree posteriors (e.g., from MrBayes) and then sampling from those quartet CF credible intervals to generate bootstrap replicates, which can then be used to estimate branch supports to annotate a candidate network. As a method that uses nucleotide sequence data as input, PhyNEST can provide a measure of branch support through bootstrap replicates. However, PhyNEST does not currently generate bootstrap replicates itself, so we did not use bootstrapping to measure branch supports for the PhyNEST networks inferred in this study. Because of the current issues with missing data in PhyNEST, the current lack of branch supports in PhyNEST compared to SNaQ, and the fact the

SNaQ network topologies were more concordant with the species tree topologies presented in Stukel et al. (2024), with a few exceptions we consider the SNaQ networks with high bootstrap support to be more indicative of the true hybridization events than the PhyNEST networks.

The D-statistic results from this study must be interpreted with caution. The D-statistic detects introgression through calculating the ratio of ABBA and BABA site patterns, which represent the alternative topologies for a gene tree of four taxa. The D-statistic assumes equal substitution rates, unlinked loci, and that the sites are a representative sample of the entire genome (Green et al. 2010; Durand et al. 2011). Our data consist of several hundred conserved nuclear genes with flanking regions, and optionally mitochondrial genomes with higher substitution rate than the nuclear genes, which violates the unlinked loci and equal substitution rates assumptions, and may not be a representative sample of the entire genome. Simulation studies have shown that violations of the equal substitution rate assumption lead the D-statistic and other summary methods to infer spurious gene flow events (Blair and Ané 2020; Frankel and Ané 2023). Our D-statistic results show many four-taxon sets with significantly unequal ABBA-BABA ratios besides those tested for our hypotheses, potentially indicating false positives.

### Reticulate evolution in Kikihia and Maoricicada

We interpret the results of the various taxon subsets of Hypotheses 1 and 2 (Westlandica Group and *K. angusta* + “murihikua” hypotheses) as evidence for the hybridization as the cause of the mitochondrial *Kikihia* “Westlandica Group” (Marshall et al. 2008, 2011; Banker et al. 2017; Stukel et al. 2024). There appears to have been several rounds of hybridization and mitochondrial capture between the species of the Westlandica Group (Figure 6). Interestingly, there is evidence of intermediate song phenotypes between species pairs involving all of the extant Westlandica Group species except for *K. subalpina* and *K.* “flemingi” (Marshall et al. 2011). This suggests that the hybridization event involving *K. subalpina* + “flemingi” was completed in the past, while hybridization among the other five species is ongoing. All of these species are restricted to the South Island of NZ except for *K. subalpina*, which is a North Island species. If the ancestor of *K. subalpina* and *K.* “flemingi” was the donor for hybridization, as suggested by the mitochondrial genome phylogeny inferred by Stukel et al. (2024), the ancestor of *K. subalpina* and *K.* “flemingi” may have been a South Island species; alternatively, the hybridization could have occurred at a time when there was still gene flow between *K. subalpina* and *K.* “flemingi” across the Cook Strait during periods of glacial maxima (Marshall et al. 2009). The other South Island Westlandica Group species currently have parapatric geographic ranges while being sympatric with *K.* “flemingi” (Marshall et al. 2008), which facilitates the current hybridization at contact zones. However, paleoclimate reconstructions show that NZ habitat types have shifted considerably since the last glacial maximum (LGM) (Alloway et al. 2007), indicating that contact zones among species were not always in their present locations.

**Figure 6:**
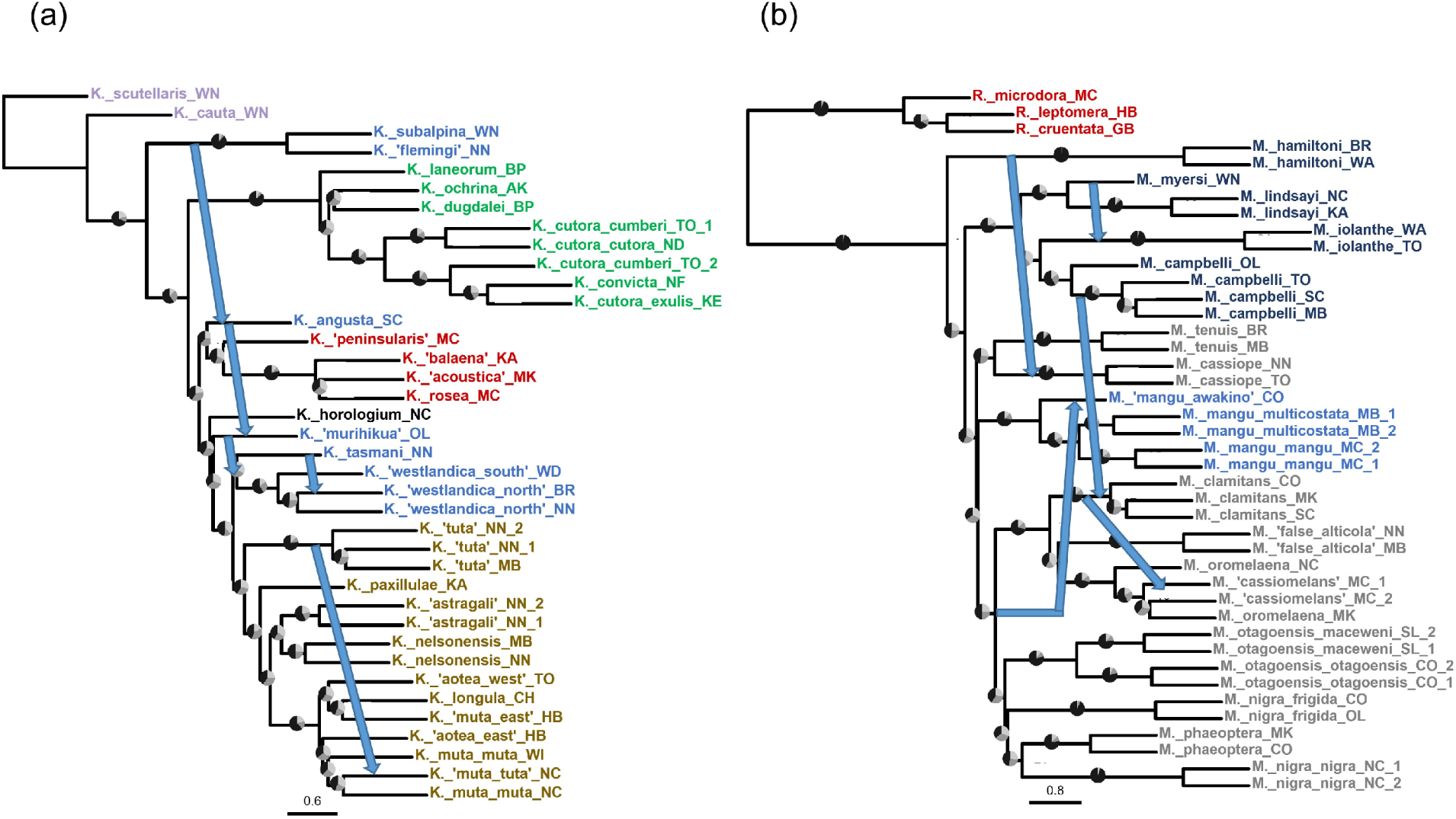
Putative hybridization events supported from the phylogenetic network and D-statistic results drawn on ASTRAL phylogeny modified from Stukel et al. (2024). (a): *Kikihia* hybridization events; (b): *Maoricicada* hybridization events.

Reconstructions of LGM species ranges for *K.* “westlandica north” and *K.* “westlandica south” have been performed (Wade 2014), but additional work needs to be done on the other species in this group to identify the likely Pleistocene contact zones.

There is interesting disagreement among the results for hybridization involving *K.* “muta-tuta”. Previous studies examining song characteristics, mitochondrial DNA, and microsatellites have demonstrated rampant, ongoing hybridization between *K. muta muta* and *K.* “tuta”, leading to capture of *K.* “tuta” mitochondrial DNA in *K. muta muta* populations – the so called *K.* “muta-tuta” individuals (Marshall et al. 2011; Wade 2014). While the D-statisitic results provide evidence of gene flow (but see discussion above on this measure), the phylogenetic network methods are inconsistent in reconstructing *K.* “muta-tuta” as a hybrid of *K. muta muta* and *K.* “tuta”. SNaQ consistently reconstructs *K.* “tuta” as the hybrid across bootstrap replicates, but as an unrooted network method it is known to occasionally have problems identifying the direction of hybrid edges (see discussion above). However, it only recovers hybridization involving *K.* “muta-tuta” with very low bootstrap support. Of the PhyNEST results, only the analysis without mitochondrial DNA was able to identify *K.* “muta-tuta” as a hybrid of *K. muta muta* and *K.* “tuta”, with the analysis using nuclear and mitochondrial DNA combined recovering hybridization with *K. paxillulae.* This is interesting, because *K. paxillulae* is not known to have a hybrid zone with *K.* “tuta”. However, *K. paxillulae* does have a hybrid zone with *K. muta muta*, including with populations that have *K.* “tuta” introgression, and Marshall et al. (2011) proposed “hybridization by proxy” to explain their finding of *K.* “tuta” mitochondrial DNA in some *K. paxillulae*. It is therefore possible that conflicting signal from this “hybridization by proxy” scenario is interfering with the phylogenetic network reconstruction.

The results for Hypotheses 4 (*M. campbelli* hypothesis) also indicate more complicated hybridization scenarios than predicted from mito-nuclear discordance. The different network topologies for *Maoricicada campbelli* and *M. iolanthe* across the alpine and lowland subsets do not show these two species involved together in hybridization with alpine species, as suggested by the nuclear and mitochondrial phylogenies inferred by Stukel et al. (2024). While the alpine PhyNEST results appear unreliable and biologically unrealistic, the alpine SNaQ results show hybridization between the alpine species and *M. campbelli* alone, while the lowland results from all analyses appear to show hybridization between *M. iolanthe* and the lowland species alone.

Some of these results may be an artifact of taxon sampling, as an alternative interpretation of the lowland results is that the apparent hybridization between *M. iolanthe* and *M. myersi* is actually the result of signal from *M. campbelli* hybridization with alpine species, which are not present in the lowland taxon subset. Additional testing with different taxon sampling schemes is needed to determine if this is the case. Regardless, hybridization events involving *M. campbelli* and *M. iolanthe* separately were suggested by the findings of Buckley et al. (2006) and are supported by the results here.

Our results are inconclusive on the history of hybridization involving *M. hamiltoni.* The findings from Stukel et al. (2024) suggested that *M. hamiltoni*, a lowland species that is sister to all other *Maoricicada* in the nuclear species trees, obtained mitochondrial DNA from the ancestor of all the other lowland *Maoricicada* species through introgression, explaining the relationships in the mitochondrial tree. None of our phylogenetic network results support this hypothesis, with the PhyNEST results appearing unreliable as discussed above, and the SNaQ results showing *M. hamiltoni* being a hybrid donor for the alpine species *M. cassiope*, albeit without high bootstrap support. It could be that mitochondrial relationships inferred by Stukel et al. (2024) are not the result of hybridization, but from ILS or some other process.

The inferences we can draw from the results of Hypotheses 6 and 7 (*M.* “mangu-awakino” and *M.* “cassiomelans” hypotheses) are clearer. There appears to be fairly strong support for the hybrid origin of the *M.* “mangu-awakino” population. However, the identity of the parent species that hybridized with *M. mangu* is unclear, shifting between *M. phaeoptera* and *M.* “false alticola” depending on the analysis. Stukel et al. (2024) suggested that the hybrid was an unsampled “ghost lineage” due to the low bootstrap support connecting *M.* “mangu awakino” to *M.* “false alticola” in the mitochondrial phylogeny. The network topologies presented here do not explicitly identify a ghost lineage as the source of introgression, but it has been pointed out that distinguishing between ghost introgression and introgression from sampled non-sister species is difficult and requires branch-length information in addition to topologies (Pang and Zhang 2024). The SNaQ results for the origin of *M.* “cassiomelans” suggest that it is a hybrid between *M. clamitans* and *M. oromelaena*, albeit in with the hybridization event inferred in the incorrect direction. *M.* “cassiomelans” individuals were originally nicknamed as such due to their appearance as normal *M. oromelaena*, but from a different locality than the rest of the *M. oromelaena* range and with songs that had characteristics resembling *M. cassiope* and *M. clamitans*, indicating a possible three-way hybrid. As our results do not show hybridization between *M. cassiope* and *M.* “cassiomelans”, either *M. cassiope* is not involved and the song similarities are illusory, or the three-way hybridization breaks the level-1 network requirement for current phylogenetic network methods. *M. cassiope* is involved according to the PhyNEST results, but the unrealistic network topology suggests that these results may be unreliable.

## Conclusion

By testing a range of hybridization hypotheses with the phylogenetic network methods SNaQ and PhyNEST, we find new insights into the reticulate evolutionary history of New Zealand cicadas. We also find that these network methods often disagree in their inferences, sometimes only slightly but other times considerably. We recommend that researchers study the natural history of their system and compare the results of multiple network methods when attempting to reconstruct phylogenetic networks, as the current methods have important limitations that must be taken into consideration. We look forward to future implementations of these network methods that may be able to address these shortcomings.

## Acknowledgements

We thank the documentation and resources prepared by the SNaQ and PhyNEST developers for assistance in performing the analyses in this study. We also acknowledge previous discussions on the PhyloNetworks-Users Google Group as well as personal discussions with Sungsik (Kevin) Kong for troubleshooting assistance with SNaQ and PhyNEST, respectively.

Computational resources for all analyses were provided by the University of Connecticut

Computational Biology Core. Paul Lewis, Elizabeth Jockusch, and Nick Matzke provided comments and suggestions on the manuscript. For help with field work we acknowledge the many people listed in Stukel et al. (2024), especially David Marshall and Kathy Hill.

## Data Availability

Sequence data, intermediate files generated from the data processing pipeline, and output from phylogenetic network analyses are deposited in tDryad, while scripts for data processing and analysis are available from Zenodo (http://datadryad.org/stash/share/MSubfB-NR9rOzzKWJMbnbtcK-Y3mX5YWmOZDTcGp0Jo).

